# DBSIM: A Platform of Simulation Resources for Genetic Epidemiology Studies

**DOI:** 10.1101/030353

**Authors:** Po-Ju Yao, Ren-Hua Chung

**Author notes:** Corresponding author: Ren-Hua Chung, PhD, Address: No 35, Keyan Road, Zhunan, Miaoli, 350, Taiwan, #36105.

## Abstract

Computer simulations are routinely conducted to evaluate new statistical methods, to compare the properties among different methods, and to mimic the real data in genetic epidemiology studies. Conducting simulation studies can become a complicated task as several challenges can occur, such as the selection of an appropriate simulation tool and the specification of parameters in the simulation model. Although abundant simulated data have been generated for human genetic research, currently there is no public database designed specifically as a repository for these simulated data. With the lack of such database, for similar studies, similar simulations may have been repeated, which resulted in redundant works. We created an online platform, DBSIM, for simulation data sharing and discussion of simulation techniques for human genetic studies. DBSIM has a database containing simulation scripts, simulated data, and documentations from published manuscripts, as well as a discussion forum, which provides a platform for discussion of the simulated data and exchanging simulation ideas. DBSIM will be useful in three aspects. Moreover, summary statistics such as the simulation tools that are most commonly used and datasets that are most frequently downloaded are provided. The statistics will be very informative for researchers to choose an appropriate simulation tool or select a common dataset for method comparisons. DBSIM can be accessed at http://dbsim.nhri.org.tw.

## Introduction

Computer simulations are routinely conducted in genetic epidemiology studies. For example, when a new statistical method is developed to test associations between genetic variants and a disease, it is important to evaluate the type I error rates for the method and compare the power of the method with other existing methods under different scenarios. Simulation studies are also important to evaluate the study design, such as case-control or family-based design, and to calculate the numbers of samples required to achieve a reasonable power when planning a genetic epidemiology study. Due to the complicated structures in human genomes and complicated disease etiology, simulating realistic genetic variants and trait values can be challenging.

A group consisting of population geneticists, genetic epidemiologists, and computational scientists addressed several current and emerging challenges and opportunities in genetic simulation studies in the “Genetic Simulation Tools for Post-Genome Wide Association Studies of Complex Diseases” workshop held at the National Institutes of Health (NIH) in Bethesda, Maryland on March 11-12, 2014 (Chen, et al., 2015). One of the challenges is that researchers may have difficulties in choosing an appropriate simulation tool from a large number of existing tools. Due to the difficulties, the researchers ended up with developing their own tools with functions overlapping those in the existing tools, which resulted in redundant works (Peng, et al., 2015). Another challenge is that simulated data for a certain study may be generated in favor of the assumptions for the statistical models developed in the study. This could lead to unfair comparisons of the method with other methods. The opportunities discussed by the group included the creation of a server for sharing genetic simulation data, identification of common datasets for method comparison, and encouragement of making simulated datasets publicly available.

In response to the challenges and opportunities addressed above, we created a platform for simulated data, DBSIM. The platform consists of a multi-functional website with user-friendly interfaces, an FTP server, and a database server. The platform was designed as a repository for simulated datasets generated from published papers or papers under peer review related to genetic epidemiology studies. DBSIM has two important features. One is that each dataset on DBSIM can be voted by the user and the other is that summary statistics, such as the datasets with the most votes, the most frequently downloaded datasets and the most frequently used simulation tools are reported on the main page of DBSIM. The summary statistics will be informative for users to select an appropriate simulation tool and a common dataset for method comparison.

## Methods

### Architecture of DBSIM

The hardware supporting DBSIM includes a server-level computer, equipped with an Intel XEON quad-core 2.4 GHz CPU and 96 GB of memory, where the computer is connected by a disk array (with a storage of 50 TB) and a Network Attached Storage (NAS) system with equal amount of storage to the disk array. The redundant array of independent disks (RAID) 4 technique was applied to the disk array as a backup mechanism for the data in case of disk failures. The data are weekly copied to the NAS system, which serves as a secondary backup mechanism for the data in the disk arrays. Web, FTP, and MySQL servers were set up on the computer.

### Web Server

A person who registered on DBSIM and deposited their simulated datasets into the database is referred to as the author, while a person who registered on DBSIM and downloaded the datasets from the database is referred to as the user. User-friendly web user interfaces (UIs) were created for the author and the user on DBSIM. The author uses an information form to specify the properties of the datasets. The information form includes some basic information about the datasets, such as a general description of the data (e.g., study design and types of data) as well as a more detailed survey of the datasets (e.g., the tools and scripts used to generate the data and technical notes for generating the data). Datasets such as simulated raw data, scripts, and any other related files are uploaded to DBSIM via the FTP server by the author. The user can search the datasets by data attributes (e.g., paper title, keywords, and author names) on the web UI, and then use the FTP server to download the datasets. The author can also leave comments and vote for a dataset on the web UI. A discussion forum is also hosted on the Web server. The forum provides a platform for questions and answers between the authors and users. On the main page of the web server of DBSIM, summary statistics such the most frequently downloaded datasets, most frequently used simulation tools, datasets with the most votes, and most viewed datasets are provided.

### MySQL Server

Data attributes are saved in the MySQL database, and queries sent from the Web server are processed by the MySQL server. Currently data on DBSIM come from either datasets deposited by the author or replicated datasets generated by our group. We selected papers that have been highly cited in the field of statistical genetics and followed the same procedures as described in the papers to generate the replicated datasets. Each dataset in DBSIM is assigned a unique identifier, which can be cited when the dataset with the corresponding identifier is used for other studies.

### FTP Server

The FTP server handles downloads and uploads of the data. Any users can download the data freely via the FTP server. Currently the author can upload files with a maximum size of 50 GB for a study. Given the current storage of 50 TB in the disk array, DBSIM will be able to accommodate data from approximately 1,000 studies. However, since the size of the data for a study can be significantly less than 50 GB, we expect that the actual number of studies that DBSIM can host will be greater than 1,000.

### Results

User friendly web interfaces have been created for the author to upload the data and for the user to search and download the data. Table 1 shows the entries of the information form that the author will need to fill in before uploading the data. Some entries such as the simulated data type and trait type are in the same format as those in Genetic Simulation Resources (GSR), a catalog of genetic data simulation software (Peng, et al., 2013). Note that although DBSIM aims to host simulated data from published studies, data from papers that are currently under review are also welcome to be deposited in DBSIM. This will provide opportunities for the editors or reviewers from the journal to assess the simulation scripts and data.

**Table 1.**
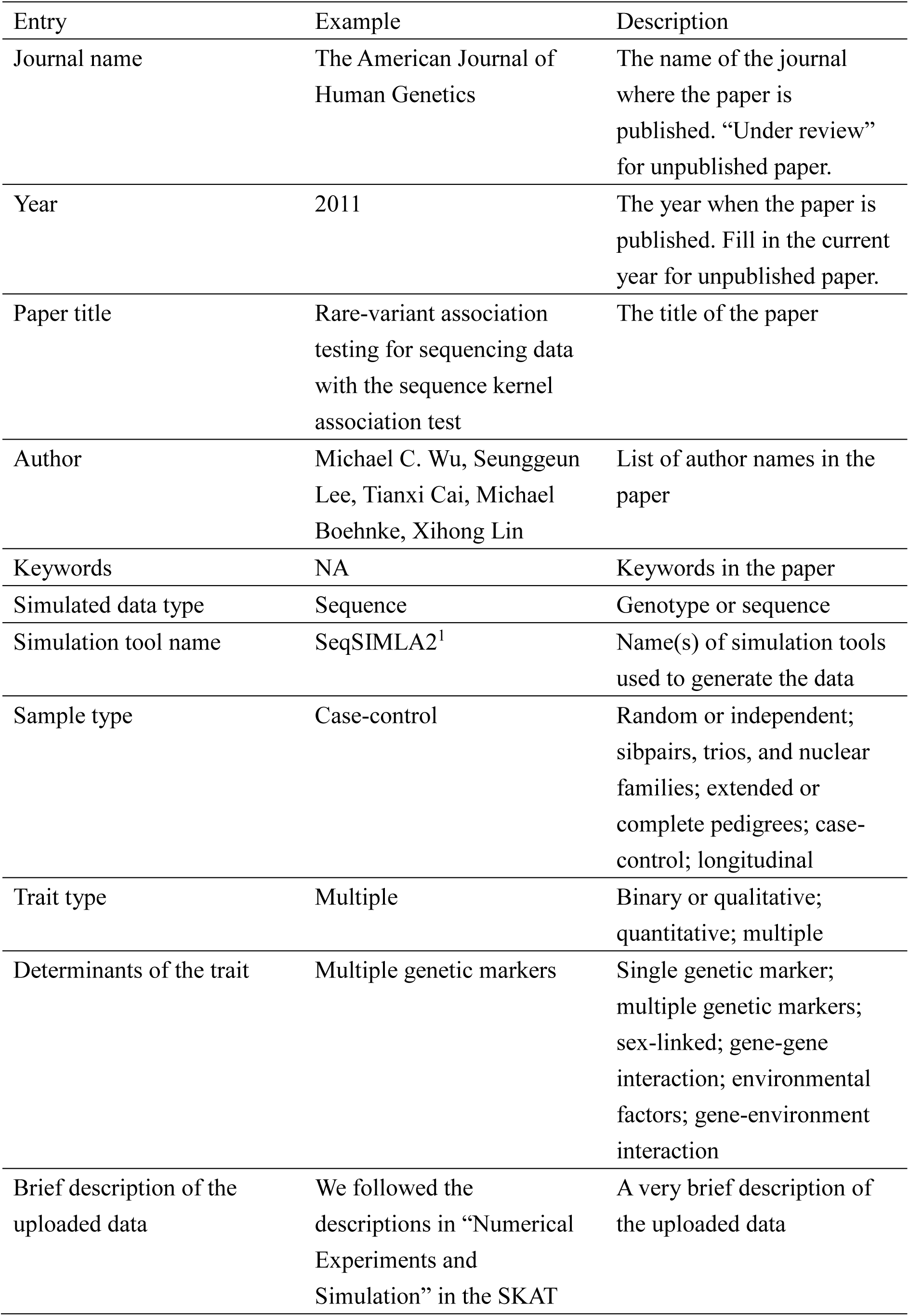
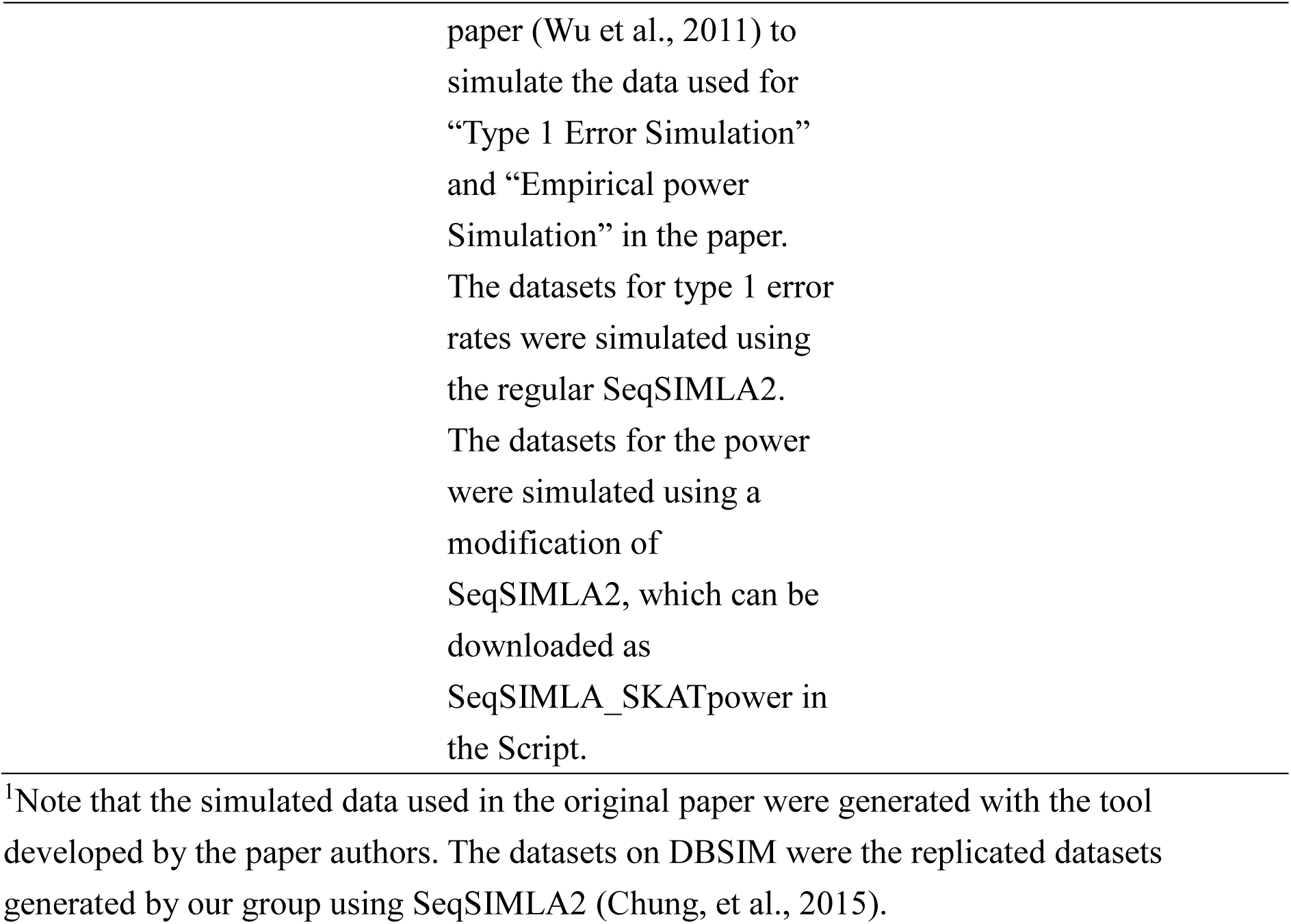
Information form for the author

Stress tests were performed for both the Web and FTP servers. Both servers functioned normally assuming 100 simultaneous users who performed regular tasks such as web browsing, searching, uploading, and downloading of the data.

The numbers of views and votes and comments from users are reported for each dataset on DBSIM. On the main page, DBSIM reports the summary statistics including the most frequently used tools, the most frequently downloaded datasets, the most viewed datasets, the datasets with the most votes, and the most viewed and voted posts in the discussion forum. The summary statistics will be informative for other simulation studies, such as choosing a simulation tool that has been widely adopted in the research community. Moreover, the most frequently downloaded datasets may become benchmark datasets for method comparisons. The forum provides an important communication platform for exchanging simulation strategies and discussion of the simulated data.

### Discussion and Conclusions

Table 2 shows the comparisons between DBSIM and two other popular public data repositories, Dryad and figshare. DBSIM has several advantages over the two repositories. DBSIM has larger storage space per study, taking into account the fact that simulated data are generally large. Moreover, several crucial summary statistics are only provided by DBSIM, such as the datasets receiving the most votes and the most frequently used simulation tools. The statistics will help eliminate the difficulties faced by the user to choose an appropriate simulation tool and help researchers to identify common datasets for method comparison. Moreover, a discussion forum is provided by DBSIM, making DBSIM not just a data repository, but also a platform for exchanging simulation strategies.

**Table 2.** Comparison between DBSIM and other public data repositories

|  | DBSIM | Dryad | figshare |
| --- | --- | --- | --- |
| Data type | Simulated data | Any | Any |
| Targeted research field | Genetic<br>epidemiology | General | General |
| Space limit | 50 GB free space<br>per study | 10 GB for about<br>\$80, and about<br>\$10 US dollars<br>for each<br>additional GB | 1 GB free space, 10<br>GB for \$8/month, 15<br>GB for \$11/month,<br>20 GB for<br>\$15/month |
| Statistics for each dataset |  |  |  |
| Number of views | Yes | Yes | Yes |
| Number of downloads | Yes | Yes | Yes |
| Number of votes<sup>1</sup> | Yes | No | No |
| Summary statistics |  |  |  |
| Most frequently<br>downloaded data | Yes | Yes | No |
| Most viewed data | Yes | No | Yes |
| Most voted data | Yes | No | No |
| Most frequently used<br>tools | Yes | No | No |
| User comment<sup>2</sup> | Yes | No | Yes |
| Share on social media | No | Yes | Yes |
| Discussion forum | Yes | No | No |
| Unique identifier | Yes (DBSIM<sup>3</sup>) | Yes (DOI<sup>4</sup>) | Yes (DOI) |

<sup>1</sup>Number of votes received by users
<sup>2</sup>Whether users can leave comments on the dataset
<sup>3</sup>The identifier is self-defined by DBSIM
<sup>4</sup>Digital Object Identifier

In conclusion, a very useful platform DBSIM was created for genetic data simulations. DBSIM will promote simulated data sharing for genetic epidemiology studies. With the information provided by DBSIM, it will become straightforward to identify the most useful and popular simulation tool, which will guide the user for selecting an appropriate tool. Also benchmark datasets can be selected, which can become common datasets for method comparisons. DBSIM aims to promote simulated data sharing and improve transparency and efficiency in simulation studies for genetic epidemiology.

## Acknowledgement

This work was funded by a grant from the National Health Research Institutes (PH-104-PP-10).

